# HLA-B51 induces IFN-*γ* production in human natural killer cells

**DOI:** 10.64898/2026.05.02.722370

**Authors:** Yasuhiro Omata, Hiroko Hayakawa, Kojiro Sato

**Affiliations:** Division of Rheumatology and Clinical Immunology, Department of Medicine, Jichi Medical University, 3311-1 Yakushiji, Shimotsuke, Tochigi 329-0498, Japan; Division of Biochemistry, Department of Medicine, Jichi Medical University, 3311-1 Yakushiji, Shimotsuke, Tochigi 329-0498, Japan

**Keywords:** Behcet’s disease, Human leukocyte antigen-B51, Human natural killer cells, KLRK1, Interferon-γ

## Abstract

Behcet’s disease (BD) is a systemic inflammatory disease. It is considered as an autoinflammatory disease triggered by innate immunity rather than adaptive immunity. Human leukocyte antigen-B51 (HLA-B51) is the strongest genetic factor associated with BD. This study investigated how HLA class 1 molecules interact with innate immune cells and induce cytokine secretion. For this purpose, 293T cells transfected with a plasmid encoding HLA-B51 were cultured with natural killer (NK) cells obtained from healthy human donors. Within 24 h, the concentrations of interleukin-4 (IL-4), IL-8, and interferon-γ (IFN-γ) in the medium increased, indicating that NK cells secreted cytokines without undergoing cellular expansion for cytolysis. NK cells stimulated by nonself HLA-B51 produced IFN-γ levels comparable to those produced by NK cells stimulated by self HLA-B51. NK cells carrying HLA-B51 were accurately recognized by overexpressing HLA-B51 on 293T cells. Moreover, ample intracellular IFN-γ levels were detected in NK cells after stimulation with phorbol 12-myristate-13-acetate (PMA) plus ionomycin. KLRK1 (CD314)-positive cells mainly primarily accounted for IFN-γ-producing cells, whereas KLRK1-negative cells did not. In contrast, both NCR1 (CD335)-positive and -negative cells contributed to IFN-γ production. We next investigated whether HLA-B51 on the surface of 293T cells stimulates KLRK1 as a ligand causing IFN-γ secretion. In masking experiments using anti-KLRK1 antibodies, NK cells with high levels of cell surface KLRK1 decreased the production of IFN-γ. Conversely, human NK cell line KHYG1 cells also produced IFN-γ in culture with 293T cells, but did not increase IFN-γ through HLA-B51 stimulation. The mRNA expression of the signal adaptor protein HCST (DAP10) in KHYG1 cells was lower than that in NK cells, whereas the relative expression of IL-2RA in KHYG1 cells was higher than that in NK cells. These findings suggest that HLA-B51 can interact with KLRK1 on the NK cells inducing IFN-γ secretion, whereas IL-2 signals outweigh HLA-51 stimulation in KHYG1 cells.

## Introduction

Behcet’s disease (BD) is a systemic inflammatory disease characterized by uveitis, vasculitis, mucocutaneous, articular, gastrointestinal, genital, and oral manifestations (1, 2, 3). BD has been categorized as an autoinflammatory disease triggered by innate immunity rather than adaptive immunity because no autoantibody has been identified. Serological studies have demonstrated that BD correlates with human leukocyte antigen-B51 (HLA-B51) (4). Moreover, genotyping studies revealed disequilibrium with MICA (5–8). Correlation with HLA class 1 alleles has been detected in other inflammatory diseases. For instance, ankylosing spondylitis, uveitis, Takayasu’s arteritis, ulcerative colitis, and psoriasis are associated with HLA class 1 molecules such as HLA-B27 (9, 10), HLA-B52 (11, 12), and HLA-C*0602 (13). Hence, a unifying concept, major histocompatibility complex (MHC)-I-opathy, has been proposed (14).

Hyperactivation of neutrophils and their invasion into the lesion are a major part of immunopathogenesis. The frequency of activated neutrophils marked by CD64 in relapse is higher than that in remission (15). Although neutrophils generally eliminate pathogens through phagocytosis, and neutrophil extracellular traps (NETs) capture pathogens, an increased formation of NETs has been detected in patients with BD with vasculitis and thrombosis (16–18).

The sterile-needle-induced damage test demonstrates γδT cells as an immediate effector. The cells are also found in the mucosal tissues of patients BD and produce interleukin-4 (IL-4) or interferon-γ (IFN-γ), thus affecting on T helper 1 (Th1) and Th2 balance (19). Cross-reaction of heat shock protein activates γδT cells and releases tumor necrosis factor-α (TNF-α), thereby recruiting neutrophils (20).

Natural killer (NK) cells exhibit surveillance function to identify nonself, infected cells, and cancer cells and produce cytokines to activate other immune cells. HLA class 1 molecules, which are expressed on cell surface where α-chain couples with β2-microglobulin (21), and their receptors on NK cells are involved in these events. An inhibitory receptor, KIR3DL1, prevents cytolysis from normal cells by identifying their own HLA-B (22, 23), whereas KIR3DL1-positive cells obtained from donors with Bw4 increase the production of IFN-γ and cause cytolysis against HLA-deficient cancer cells (23, 24). Regarding cytokine production, CD56^bright^ NK cells represent cytokine-producing cells, and are found from early to middle stages of the differentiation step. In BD, despite the decreased numbers of both CD56^bright^ and CD56^dim^ NK cells in peripheral blood, CD56^bright^ NK cells obtained from patients with BD contain more IFN-γ-producing cells than those obtained from healthy donors (25). Considering cytokine production, NK cells are classified into two groups based on IFN-γ production (26). NK1 cells produce IFN-γ, TNF-α, and IL-2, whereas NK2 cells produce IL-4, IL-5, and IL-13. Increased NK1/NK2 ratios have been detected in patients with uveitis and mucocutaneous BD (27, 28). Moreover, single-cell analysis identified the expansion of C1q^high^ monocytes in patients with BD (29), which provides a prospect that IFN-γ had already activated the secretion of proinflammatory cytokines before peripheral blood was sampled from patients.

Several receptors mediate stimulation. For instance, KLRK1 (CD314) is expressed on the surface of almost all NK cells. With the adaptor protein HCST (DAP10), KLRK1 transmits activating signals and results in cytolysis (30, 31). There are also several ligands that activate KLRK1 and induce different reactions. For instance, one of the MHC class 1-related chain, MICA, activates KLRK1 and induces cytolysis (32). Moreover, ULBPs also bind to KLRK1, inducing the production of cytokines (33, 34). NCR1 (CD335) eliminates cancer cells with low levels of HLA class 1 molecules (35). Nevertheless, there is limited information regarding the roles of HLA class 1 molecules in immediate cytokine production.

In this study, we developed an *in vitro* coculture of 293T cells with human NK cells to investigate how HLA-B51 activates NK cells and induces cytokine secretion. HLA-B51 on the surface of 293T cells induced IFN-γ production in NK cells. We also demonstrated that HLA-B51 binds to KLRK1 on the surface of NK cells. In contrast, KHYG1 cells produced IFN-γ with IL-2 input mediated through IL-2 receptors.

## Materials and methods

### Cell culture

293T cells and KHYG1 cells were obtained from KAC Co., Ltd. (Kyoto, Japan) and JCRB Cell Bank (Osaka, Japan), respectively. 293T cells were maintained in DMEM (4.5 g/L glucose, Nacalai Tesque, Kyoto, Japan) supplemented with 10% fetal bovine serum (Sigma-Aldrich Inc. St. Louis, MO, USA) and penicillin-streptomycin solution (×50; FUJIFILM Wako Pure Chemical, Osaka, Japan) at 37°C in a humidified chamber containing 5% CO_2_. KHYG1 cells were maintained in RPMI 1640 (Nacalai Tesque) supplemented with 10% fetal bovine serum (Sigma-Aldrich Inc.), penicillin-streptomycin solution (×50), 100 mM sodium pyruvate solution (Nacalai Tesque), MEM nonessential amino acids solution (×100; Nacalai Tesque), and IL-2 (20 ng/mL, Peprotech, Cranbury, NJ, USA) at 37°C in a humidified chamber containing 5% CO_2_.

### Isolation of human NK cells

From healthy donors, 20 mL of peripheral blood was collected with heparin (MOCHIDA PHARMACEUTICAL CO. LTD, Tokyo, Japan), and was mixed with the same amount of phosphate-buffered saline (PBS). The diluted blood was layered on 10 mL of Ficoll-Paque™ PLUS (Cytiva, Tokyo, Japan). After centrifugation at 1,800 rpm for 15 min at 27°C, the mononuclear cell layer was collected, suspended in PBS, and centrifuged again at 1,600 rpm for 5 min. Pellets were washed with PBS again by centrifugation at 1,400 rpm for 5 min and resuspended with DMEM (Nacalai Tesque). NK cells were separated from peripheral blood mononuclear cells (PBMCs) using the autoMACS system with human NK Cell Isolation Kit (Miltenyi Biotec, Bergisch Gladbach, Germany) according to the manufacturer’s procedures. HLA class 1 types were examined using antigen tests. This study was approved by the Ethics Review Committee for University Clinical Research of Jichi Medical University (20-148).

### Coculture of 293T and NK cells

293T cells (3.6 × 10^4^ cells) were cultured on Falcon 48-well plates (Corning Inc., Corning, NY, USA). Plasmid DNA (Riken BRC DNA BANK, Tsukuba, Japan) was mixed with FuGENE HD Transfection Reagent (Promega, Madison, WI, USA) and Opti-MEM™ Reduced Serum Medium (Thermo Fischer Scientific, Waltham, MA, USA), and then transfected into 293T cells. After 24 h, a suspension of NK cells (2.5 × 10^5^ cells) or KHYG1 cells (2.5 × 10^5^ cells) was added to the culture. For KLRK1 blocking, NK cells were incubated with anti-human NKG2D/CD314 antibody (2.5 µg; clone #149810, R&D SYSTEMS, Minneapolis, MN, USA) for 10 min before adding to the coculture. Anti-mouse IgG1 antibody (2.5 µg; clone #11711, R&D SYSTEMS) was used as a control.

Plasmid DNA was handled according to the Ethic Review Committee for Gene Recombination of Jichi Medical University (R5-07).

### Cell stimulation

NK cells and KHYG1 cells were incubated with a pre-mixed solution of phorbol 12-myristate-13-acetate (PMA) plus ionomycin (Cell Activation Cocktail without Brefeldin A, BioLegend, San Diego, California, USA) and GolgiStop (BD Biosciences, Franklin Lakes, NJ, USA) for 4 h. The cells were washed with PBS for the detection of intracellular IFN-γ. Culture media collected from cells treated with PMA plus ionomycin were analyzed by ELISA.

### Bead array

Diluted supernatant was analyzed using Bio-Plex Suspension Array System (BioRad, Hercules, CA, USA) with Human Cytokine 17-plex Panel (BioRad) according to the manufacturer’s protocol.

### ELISA

The culture media collected from the coculture or NK cells stimulated with PMA plus ionomycin were centrifuged for 1 min, and the resulting supernatant was analyzed. The concentration of IFN-γ was determined using Human IFN-γ Quantikine ELISA kit (R&D SYSTEMS) and a microplate reader (TECAN, Männedorf, canton of Zürich, Switzerland)

### qRT-PCR

qRT-PCR analysis was performed as described previously (36). Briefly, total RNA was extracted from 293T cells, NK cells and KHYG1 cells using the RNeasy Micro Kit (Qiagen, Venlo, The Netherlands). The concentration of total RNA was measured using a NanoDrop One (Thermo Fisher Scientific). Next, cDNA was synthesized from the total RNA (200 ng) using the PrimeScript™ II 1^st^ strand cDNA Synthesis Kit (Takara Bio Inc. Kusatsu, Shiga, Japan). The cDNA (5 ng) was amplified using Power SYBR™ Green PCR Master Mix (Thermo Fisher Scientific) in StepOnePlus Real-Time PCR (Applied Biosystems, Waltham, MA, USA). The relative expression of mRNA was quantified using the ΔΔCt method. The primer sequences designed for SYBR Green detection are listed in Table 1.

**Table 1.**
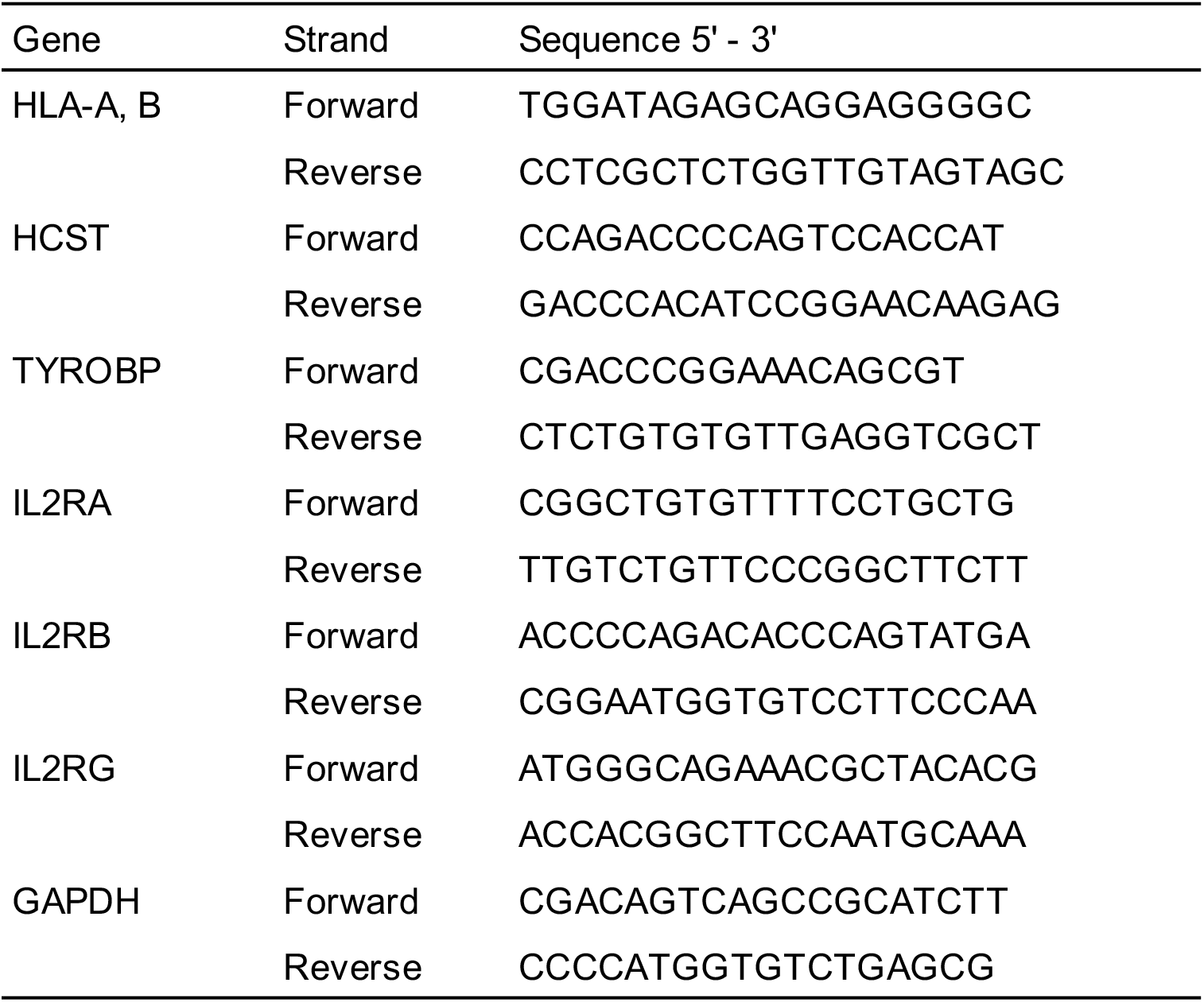
qRT-PCR primer sequences.

### KIR typing

Genomic DNA was extracted from NK cells, KHYG1 cells, and 293T cells using NucleoSpin Blood (Takara). The DNA (10 ng) was amplified using KOD FX (Toyobo, Osaka, Japan) with the following cycling program: one cycle at 94 °C for 120 s, followed by 40 cycles at 98 °C for 10 s, 65 °C for 30 s, and 68 °C for 30 s. The PCR products were confirmed by agarose gel electrophoresis. The primer sequences described by Jones DC et al. (2006) were used (37).

### Western blotting

Western blotting was conducted as described previously (36). Briefly, 293T cells were lysed in RIPA buffer (Nacalai Tesque). Protein concentration was determined using a bicinchoninic acid (BCA) assay (Thermo Fisher Scientific). The denatured lysate mixed with Novex™ Tris-Glycine SDS Sample Buffer (2×; Thermo Fisher Scientific) was loaded onto Novex™ WedgeWell™ 8%–16% Tris-Glycine gels (Thermo Fisher Scientific) for electrophoresis. The resulting gel was electroblotted onto a PVDF membrane using iBlot 2 PVDF mini stacks (Thermo Fisher Scientific). The membrane was blocked with 5% skim milk (FUJIFILM Wako Pure Chemical Corporation), and then incubated overnight with primary antibody at 4°C, followed by incubation with secondary antibody at 27°C for 1 h. Next, ECL Prime Western Blotting Detection Reagent (GE Healthcare Bioscience, Chicago, IL, USA) was added to the membrane, and the chemiluminescent signal was detected using a CCD camera (Vilber Bio Imaging, Collégien, France). Mouse monoclonal antibodies against HLA-ABC (Proteintech, Rosemont, IL, USA, clone 5C5B7, 1:5,000 dilution) and β-actin (Sigma-Aldrich, A1978, 1:2,000 dilution) were used as the primary antibodies. Horse radish peroxidase-conjugated anti-mouse antibodies were used as the secondary antibody (GE Healthcare Bioscience. 1:10,000 dilution).

#### Flow cytometry

Cells were incubated with Fixable Viability Stain 780 (BD Biosciences) for 15 min at 27°C and then blocked with FcR Blocking Reagent (Miltenyi Biotec) for 15 min at 4°C. Then, the cells were stained with specific antibodies for 20 min at 4°C. Finally, the cells were incubated with Fixation Buffer (BioLegend) for 20 min and washed twice with PBS. Cytofluorometric analyses were performed on a BD LSRF Fortessa X-20 flow cytometer (BD Biosciences) and data was analyzed using the FlowJo software (BD Biosciences). The following antibodies were used in this study: APC anti-human HLA class 1 Bw4, REAffinity™ (Miltenyi Biotech), PE anti-human HLA-A, B, C antibody (clone w6/32, BioLegend), Brilliant Violet 421™ anti-human β2-microglbulin antibody (A17082A, BioLegend), FITC anti-human CD335 (NKp46) antibody (clone 9E2, BioLegend), and PE anti-human CD314 (NKG2D) antibody (clone 1D11, Biolegend), As controls, mouse IgG1 k isotype control antibodies conjugated with each fluorescence were used (clone MOPC-21, Biolegend).

For intracellular staining, BD Cytofix/Cytoperm™ Fixation/Permeabilization Kit (BD Biosciences) was used. After staining for cell surface antigens, the cells were incubated with Cytofix/Permeabilization solution for 15 min at 27°C. Then, the cells were washed with Perm/Wash Buffer, followed by incubation with Brilliant Violet 421™ anti-human IFN-γ antibody (clone 4S.B3, BioLegend).

## Results

### *In vitro* model for cytokine production

To reproduce the proinflammatory cytokines induced by HLA class 1 molecules, we developed a coculture of 293T cells that express HLA-B51 with human PBMCs. Although 293T cells possess own HLA-A3 and HLA-B7 (38), an increased expression of HLA-A and HLA-B mRNA was observed 24 h after the transfection (Figure 1A). Moreover, strong levels of HLA class 1 expression were detected by anti-HLA class 1 ABC antibody in 293T cells at 24 h (Figure 1B, C). Flow cytometric analysis revealed an increase in the HLA-Bw4-positive population distinct from the endogenous HLA-A3- and HLA-B7-positive population (Figure 1D-H). The expression of HLA-B51 was also confirmed as shifted peaks on histograms (Figure 1G) and substantial increases in the mean fluorescent intensity (MFI) (Figure 1H). Endogenous β2-microglubulin formed a stable complex with HLA class 1 molecules on 293T cells (Figure S1A). Moreover, the HLA-Bw4 antibody detected the cell surface expression of HLA-B51 with β2-microglobulin (Figure S1B).

**FIGURE 1.**
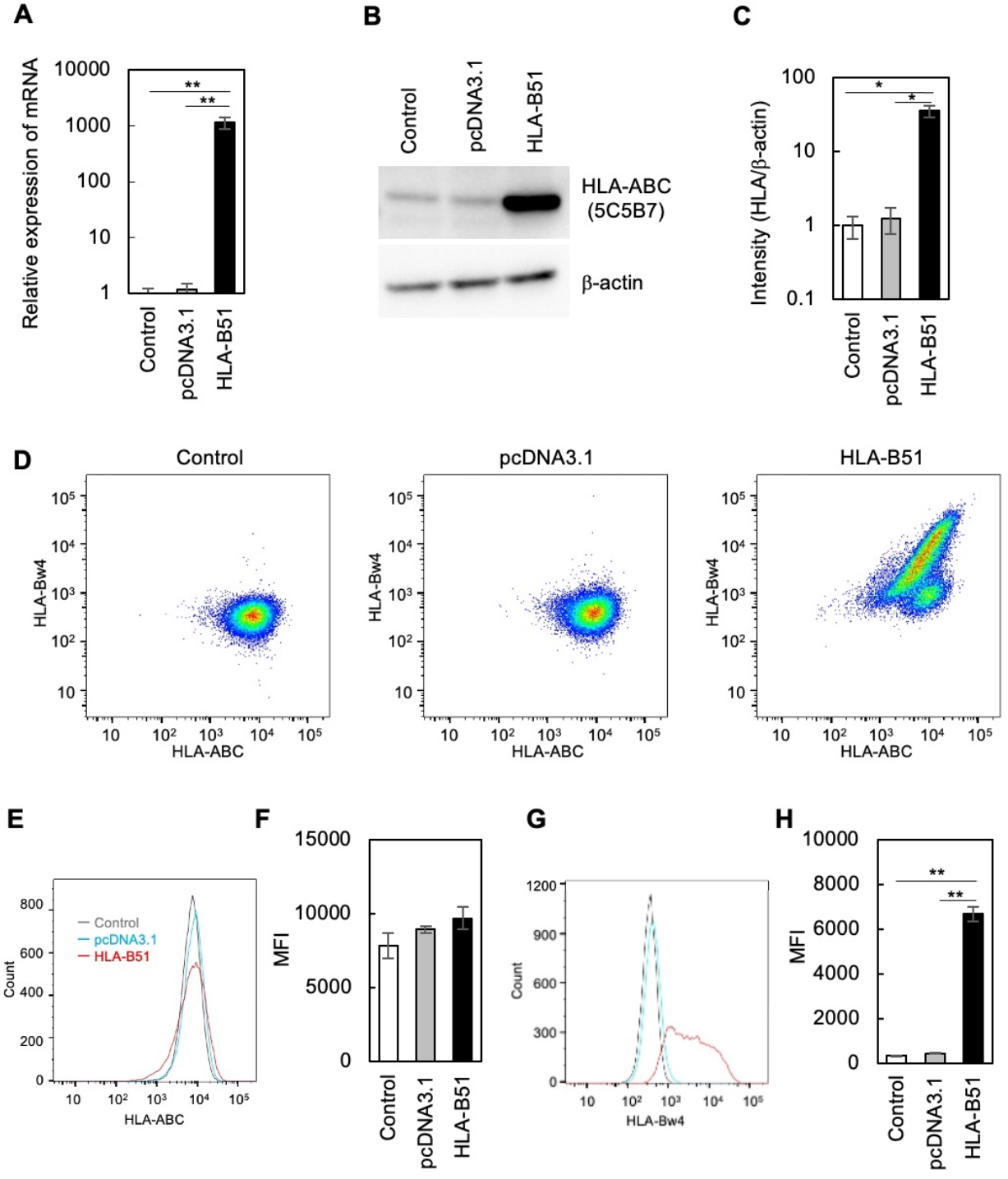
Coculture of 293T cells with NK cells for cytokine production (A) The expression of HLA-A and HLA-B in 293T cells at 24 h after transfection was analyzed by qRT-PCR. The relative expression levels were determined using the ΔCt method. Significant differences were analyzed using pairwise Wilcoxon test (** p. adjust<0.01, mean ± SE (n = 3)). (B, C) The HLA-B protein expression is depicted on a western blot. At 24 h after transfection, 293T cells were lysed and electrophoresed on an SDS-polyacrylamide gel (SDS-PAGE). Signal intensity was quantified. Significant differences were examined by pairwise *t*-test (* p. adjust<0.05, mean ± SE (n = 4)). (D-H) The cell surface expression was analyzed by flow cytometry using anti-HLA ABC and Bw4 antibodies. Representative data are shown in panel D. The surface expression is depicted as histograms (E, G) and MFI (F, H). Significant differences were examined by pairwise *t*-test (** p. adjust<0.01, mean ± SE (n = 3)).

### Cytokines secreted from NK cells

We cultured 293T cells with PBMCs and analyzed the levels of cytokines in the culture medium using bead arrays, which revealed no significant increases in the levels (data not shown). We next separated NK cells from healthy peripheral blood and cultured them with 293T cells expressing HLA-B51. Data obtained from bead arrays were subjected to clustering and heatmap analysis. Among 17 cytokines, the concentrations of IL-4, IL-8, and IFN-γ in the culture media of 293T cells expressing HLA-B51 with NK cells were higher than those in the control (Figure 2A). NK cells obtained from Donors 1 and 2 were HLA-B51-negative, whereas those from Donor 3 were positive. HLA-B51 expressed on the 293T cells induced IFN-γ production in NK cells irrespective of whether they are nonself or self (Figure 2B). The variance in IFN-γ production in HLA-B51-positive NK cells was lower than in HLA-B51-negative cells (Figure 2C). We also examined IFN-γ production from an NK cell line, KHYG1 cells, established from aggressive NK cell leukemia (39). Although KHYG1 cells produced IFN-γ in culture with 293T cells, HLA-B51 expressed on the surface of 293T did not increase the production of IFN-γ (Figure 2D).

**FIGURE 2.**
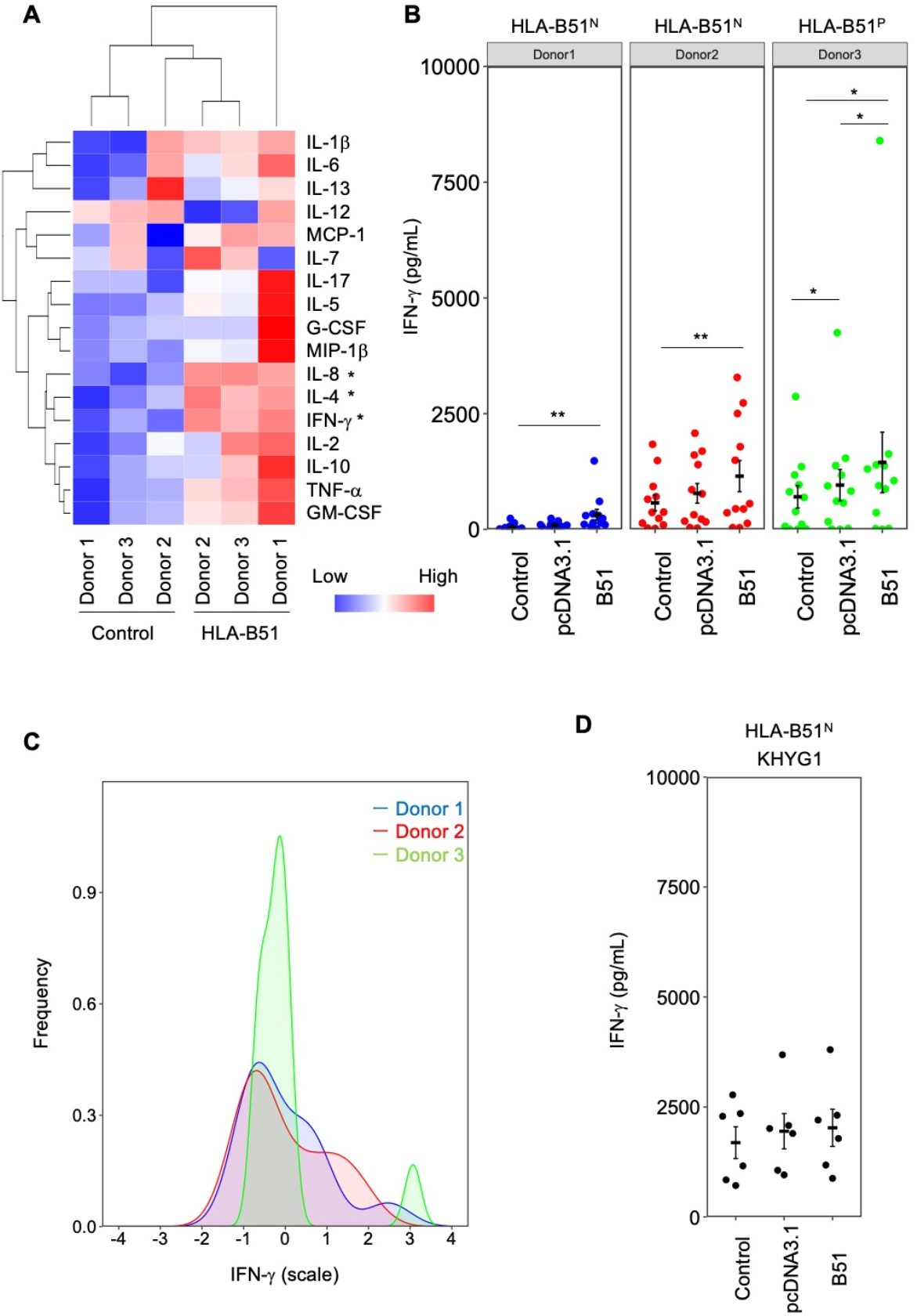
IFN-γ secretion from NK cells (A) NK cells obtained from human healthy donors were cultured for 24 h with 293T cells expressing HLA-B51. The concentration of cytokines in supernatant was analyzed on bead arrays. Normalized data were subjected to clustering and heatmap analysis. Statistical significance was analyzed using Student *t*-test (n = 3, * q <0.05). False discovery rate was controlled using the Benjamini-Hochberg procedure. (B) The concentration of IFN-γ in culture with NK cells was measured by ELISA. Statistical significances were analyzed by pairwise Wilcoxon test (** p<0.01, * p <0.05, mean ± SE (n = 12)). (C) The variance in IFN-γ production from NK cells is depicted as histograms. (D) KHYG1 cells were cultured with HLA-B51-expressing 293T cells. The concentration of IFN-γ was determined by ELISA (mean ± SE (n = 6)).

### IFN-γ secretion caused by PMA plus ionomycin stimulation

We examined the ability of NK cells and KHYG1 cells to produce IFN-γ by chemical stimulation. Intracellular staining revealed IFN-γ production in both NK cells and KHYG1 cells (Figure 3A, C). The majority of NK cells were KLRK1-positive and approximately 50.9% - 77.9% of them contained intracellular IFN-γ positive cells (Figure 3B). Regarding NCR1, a triggering receptor for cytolysis (35), the ratios of IFN-γ-positive cells in NCR1-negative cells were 41.5%–55.8%, whereas those in NCR1-positive cells were 9.3%–32.4% (Figure 3D). Both NK cells and KHYG1 cells included cytokine-producing cells, and there were no significant differences in the levels of intracellular IFN-γ between NK cells and KHYG1 cells (Figure 3E). We also examined the culture medium from NK cells and KHYG1 cells treated with PMA plus ionomycin. Treatment with GolgiStop, a protein transport inhibitor, decreased the amount of IFN-γ secretion (Figure 3F). Stimulation with PMA plus ionomycin increased IFN-γ secretion; however, IFN-γ was successfully tethered to the Golgi apparatus in NK cells and KHYG1 cells in the presence of GoligiStop.

**FIGURE 3.**
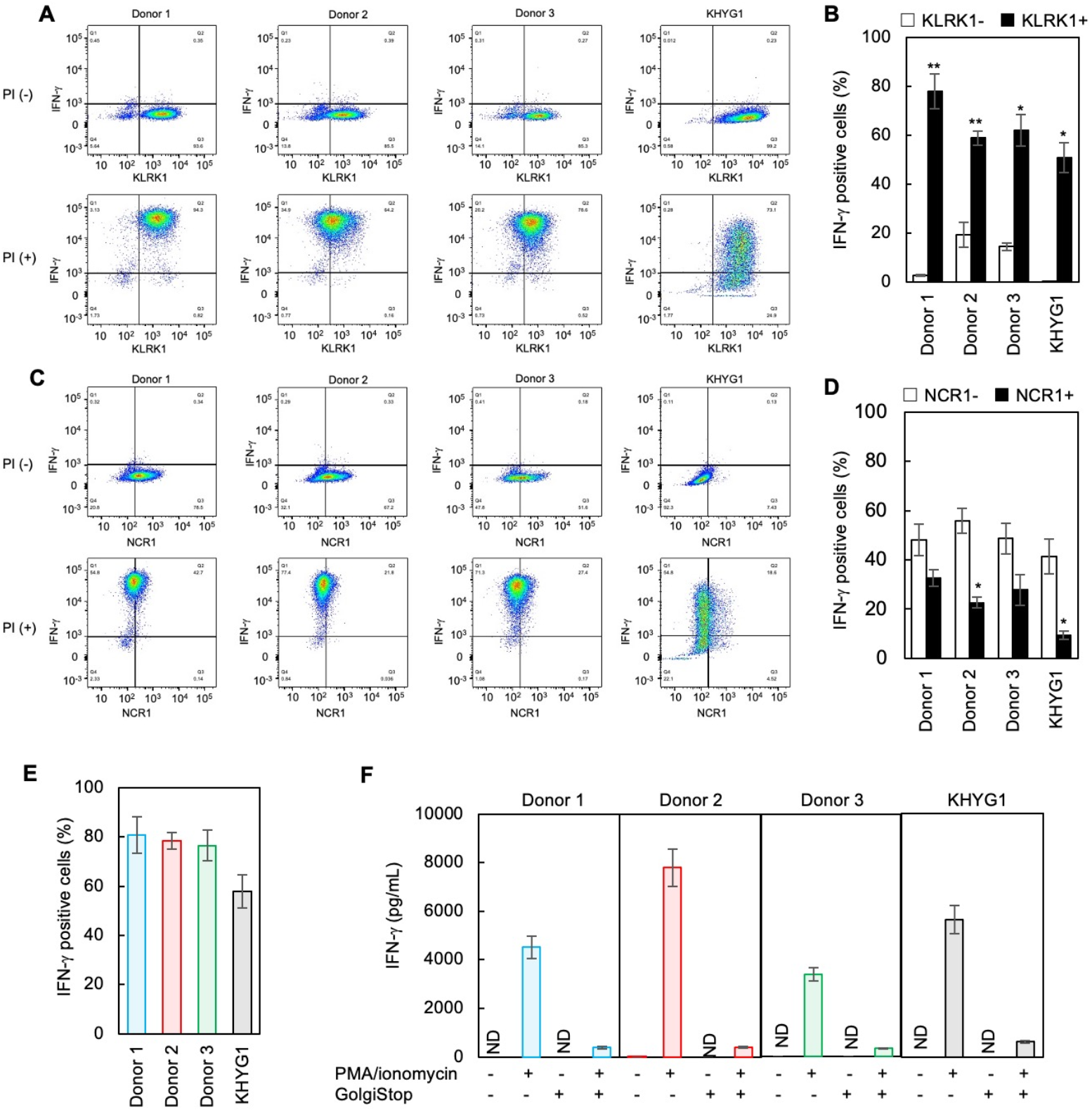
IFN-γ production in NK cells stimulated with PMA plus ionomycin (A-E) NK cells and KHYG1 cells were incubated with PMA plus ionomycin for 4 h. Cells stained with anti-human KLRK1 (CD314), anti-human NCR1 (CD335), and anti-human IFN-γ were analyzed by flow cytometry. Representative staining of intracellular IFN-γ with KLRK1 (A) or NCR1 (C), and the ratios of IFN-γ-positive cells to KLRK1-positive or -negative cells (B) and to NCR1-positive or -negative cells (D) are shown. Statistical significance was analyzed using Student’s *t*-test (** p<0.01, * p<0.05, mean ± SE (n = 3)). (E) The ratios of IFN-γ-positive cells in NK cells and KHYG1 cells were calculated. (F) The concentration of IFN-γ in the supernatant obtained from NK cells and KHYG1 cells treated with PMA plus ionomycin and/or GolgiStop is shown (mean ± SE (n = 3)).

### Interaction of between HLA-B51 and KLRK1

We examined the cell surface expression of KLRK1 and NCR1 in NK cells and KHYG1 cells were analyzed by flow cytometry (Figure 4A). NK cells exhibited lower KLRK1 MFI and higher NCR1 MFI than KHYG1 cells (Figure 4B, C). Remarkably, NK cells obtained from Donor 1 exhibited higher KLRK1 MFI than those obtained from Donors 2 and 3 (Figure 4B), whereas Donor 3 showed higher NCR1 MFI than those obtained from Donors 1 and 2 (Figure 4C). To determine whether HLA-B51 interacts with KLRK1 on the surface of NK cells, the KLRK1 receptor was blocked using anti-KLRK1 antibody. We observed decreased amount of IFN-γ secretion in the culture medium of NK cells obtained from Donor 1, but not in those obtained from other donors (Figure 4D).

**FIGURE 4.**
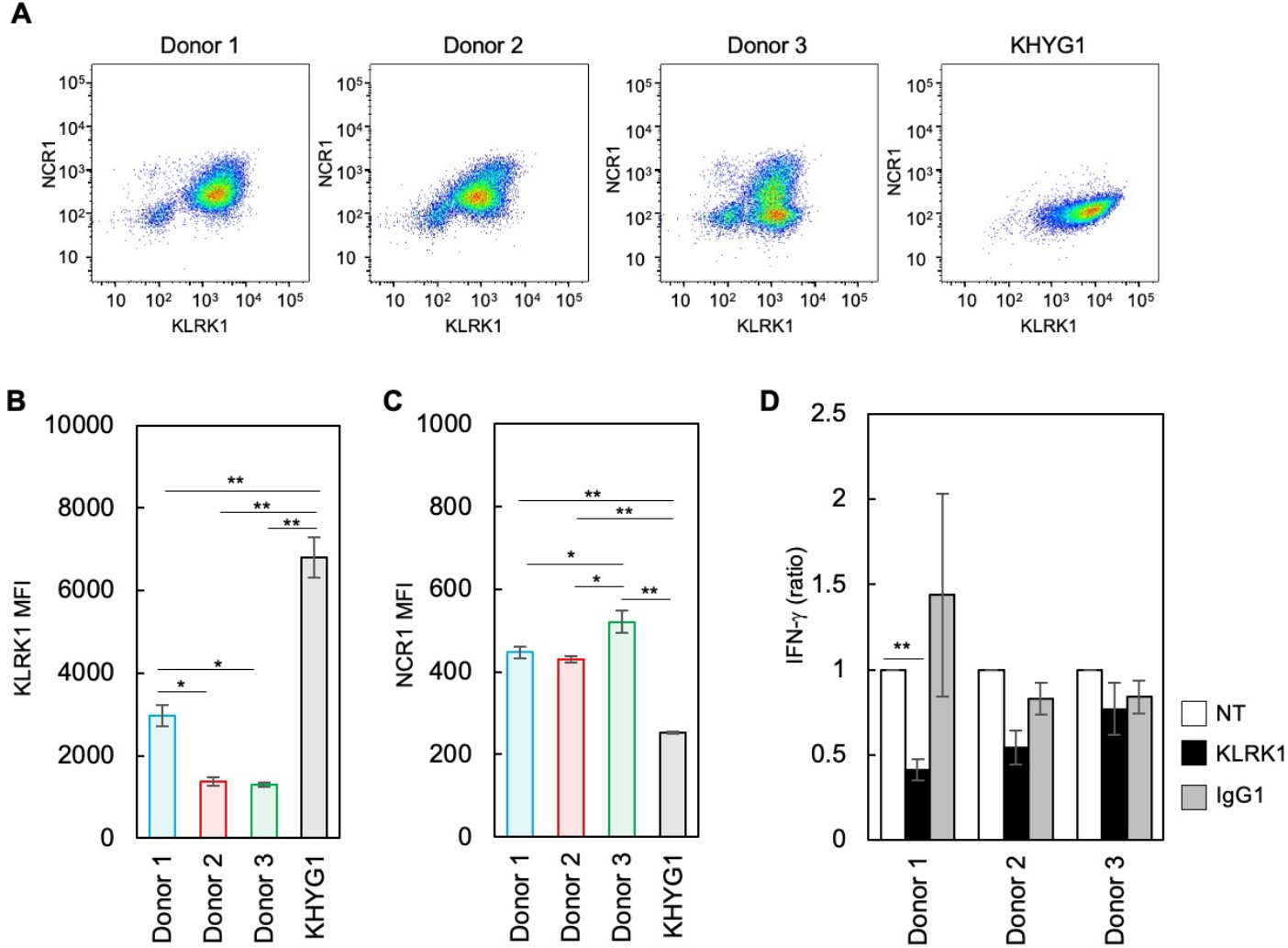
Cell surface expression of KLRK1 and its interaction with HLA-B51 (A-C) NK cells and KHYG1 cells stained with anti-human KLRK1 and anti-human NCR1, which were analyzed by flow cytometry. Representative data (A) and MFI for KLRK1 and NCR1 are depicted (B, C). Statistical significance was analyzed using the Tukey test (** p<0.01, * p<0.05, mean ± SE (n = 3)). (D) HLA-B51-expressing 293T cells and NK cells were cultured with anti-human KLRK1 for 24 h. The concentration of IFN-γ was determined by ELISA. Statistical significance was analyzed by pairwise *t*-test (** p. adjust<0.01, mean ± SE (n = 4)).

### IFN-γ production in KHYG1 cells

KHYG1 cells require IL-2 for survival and proliferation, and IL-2 induces IFN-γ production (39). To clarify whether IL-2 signals outperform the HLA-B51 stimulation on the cells, we cultured KHYG1 cells with 293T cells expressing HLA-B51 in different concentrations of IL-2 and found increased levels of IFN-γ production depending on IL-2 concentration (Figure 5A).

**FIGURE 5.**
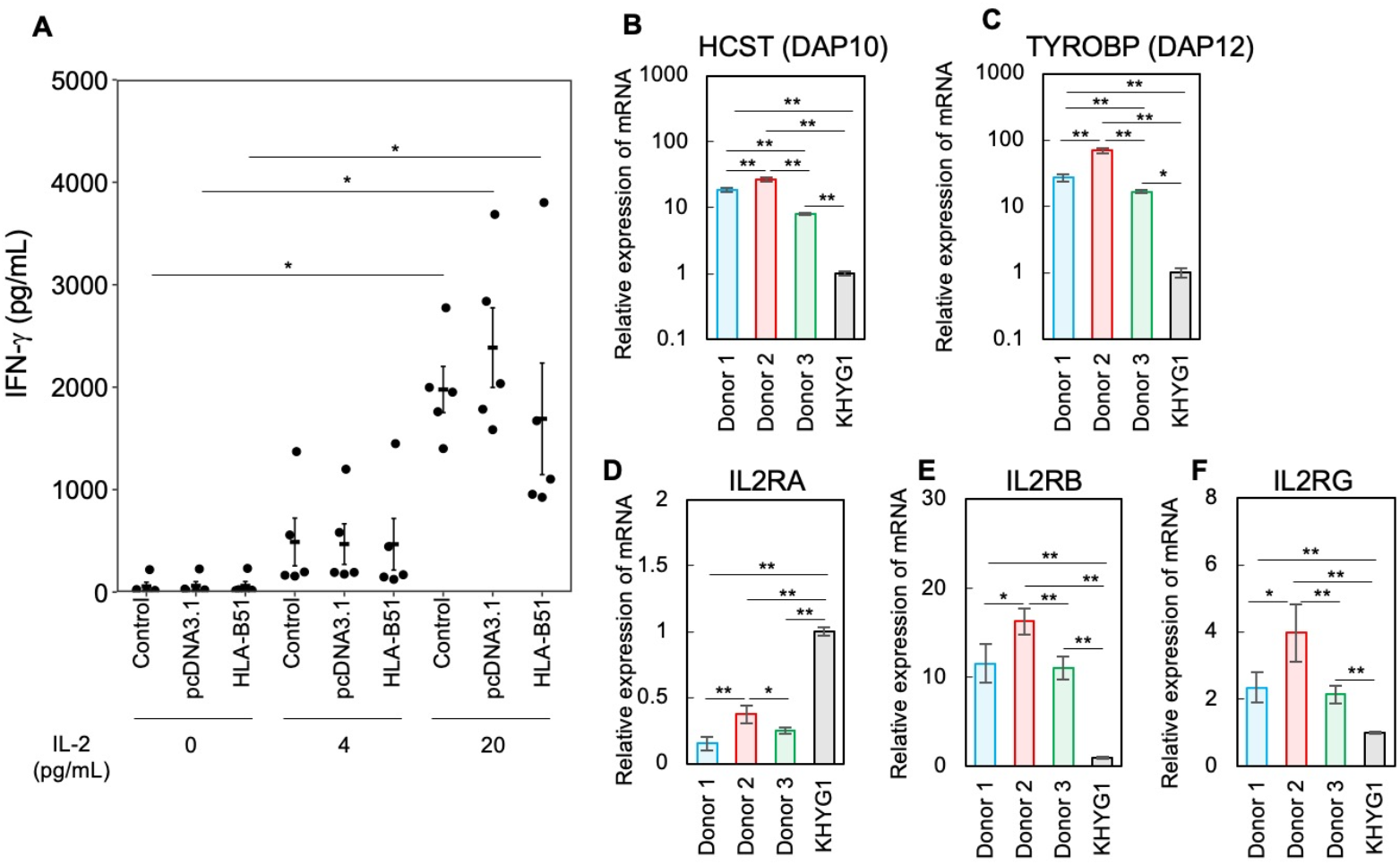
IFN-γ production in KHYG1 cells through IL-2 signaling (A) KHYG1 cells were cultured with 293T cells expressing HLA-B51 in the presence of IL-2. The concentration of IFN-γ was determined by ELISA. Significant differences were analyzed using the Steel test (* p<0.05, mean ± SE (n = 5)). (B-F) The relative expression of mRNA in NK cells and KHYG1 cells was determined by qRT-PCR. Statistical significance was analyzed using the Steel Dwass test or Tukey test (** p<0.01, * p<0.05, mean ± SE (n = 3)).

To explore the difference in IFN-γ production in NK cells and KHYG1 cells, we compared the expression of adaptors and IL-2 receptors, and found that the relative expression levels of both HCST and TYROBP in KHYG1 cells were much lower than those in NK cells (Figure 5B, C). IL-2 receptors have different affinities to IL-2. IL2RA exhibits high affinity, whereas IL2RB and IL2RG show low affinity (40). The relative expression of IL2RA in KHYG1 cells was higher than that in NK cells (Figure 5D). In contrast, the relative expression of IL2RB and IL2RG in KHYG1 cells was lower than that in NK cells (Figure 5E, F). Altogether, KHYG1 cells cannot activate intracellular signaling molecules via KLRK1 and are more responsive to IL-2 than NK cells.

## Discussion

Stimulating NK cells with PMA plus ionomycin, polyinosinic-polycytidylic acid (Poly (I:C)), antibodies against receptors, or culture with other cells has been performed for analyzing production of cytokines (41–46); however, these studies lack ligands input mediated through receptors. In the present study, we developed a coculture of 293T cells with NK cells, where HLA-B51 stimulates NK cells, resulting in IFN-γ production (Figure 2B). The production of IFN-γ in NK cells indicates inflammatory cascades in BD, where IFN-γ activates a subpopulation of macrophages (29). Although it is difficult to completely exclude a counterargument that NK cells obtained from healthy donors were infected with a virus and affected IFN-γ production, at least NK cells are capable of secreting IFN-γ via HLA-B51 stimulation on the surface of virus-free 293T cells. Meanwhile, IL-8 activates neutrophil chemotaxis and NET formation in BD (47, 48). IL-8 production in the coculture (Figure 2A) was consistent with increased IL-8 levels in patients with active BD (49).

The coculture allowed HLA-B51 to interact with NK receptors and induce immediate cytokine production. A cytolytic potential in NK cell lines that release IFN-γ with perforin and granzymes (50, 51) raises a concern that they caused cytolysis in the culture with 293T cells. Nevertheless, the mechanism underlying the immediate IFN-γ secretion and cytolytic IFN-γ is fundamentally different considering that recycling endosome (RE) orchestrates IFN-γ secretion as a post-Golgi component, whereas cytotoxic granules do not require RE (52). Another reason is that unlike K562 cells whose HLA class 1 molecules are masked with the antibody-restoring susceptibility to cytolysis (53), 293T cells that possess increased levels of HLA-B51 were less likely to be lysed. Furthermore, NK cells can acquire improved cytotoxicity after they undergo the expansion and activation process on feeder cells (54). Therefore, we argue that IFN-γ production within 24 h induced by HLA-B51 stimulation is distinct from IFN-γ release in cytolysis. Interestingly, NK cells that exhibited increases in cytokine production against 293T cells expressing high levels of HLA-B51 were derived from KIR3DL1-positive donors (Figure 2B, Figure S2). KIR3DL1 could recognize HLA-B51 and produce IFN-γ; however, KIR3DL1-positive cells are not a major population of peripheral NK cells (22). Masking experiments revealed that HLA-B51 recognized the high levels of KLRK1 (Donor 1) (Figure 4B, D). The affinity of HLA-B51 for KLRK1 might not be strong, and HLA-B51 could pair with other receptors when the levels of KLRK1 are low. Heterodimers of KLRD1 and KLRC2 can pair with TYROBP that contains an immunoreceptor tyrosine-based activation motif and activate NK cells (55), whereas HLA-B7, which is classified as Bw6, recognizes KLRD1 in target cell lysis (56).

The different levels of cell surface KLRK1 from healthy donors (Figure 4B) raised the question how the transcription of KLRK1 is regulated. A hypothesis is that splicing variants form different structures on the cell surface. The full length of KLRK1 has a ligand-binding ectodomain that is detectable in flow cytometry; however, variants whose introns are not eliminated lack the ectodomain and result in exhibiting low levels of surface expression. Because these truncated variants bind HCST, they interfere with the interaction of full-length KLRK1 with HCST (57). Another possibility is the KLRC4– KLRK1 readthrough gene. KLRK1 is located just next to KLRC4 on chromosome 12, both of which possess C-type lectin domain (58). KLRC4 does not form extracellular structures despite the ability to pair with TYROBP (59). Because the transcription machinery cannot identify a termination site (60), KLRC4–KLRK1 could generate hybrid proteins. Alternatively, it might include different lengths of protein using several promoters and transcription initiation sites. Certain types of KLRC4-KLRK1 have the potential to form extracellular structures that could be activated by HLA-B51.

NCR1 is a triggering receptor on NK cells that induces cytotoxicity, and NCR1^bright^ cells have been identified as the population that lyses cancer cells (35). The presence of intracellular IFN-γ in NCR1 negative cells (Figure 3D) supports the idea that NK cells and KHYG1 cells include cytokine-producing cells.

Although IL-10 increases IFN-γ production in NK cells with IL-18 (61), we detected IFN-γ in the culture medium of NK cells through HLA-B51 stimulation without IL-10 and IL-18 (Figure 2A, B). Moreover, treatment with PMA plus ionomycin treatment produced almost equal levels of IFN-γ in NK cells (Figure 3E). PMA activates protein kinase C (PKC), whereas ionomycin, a calcium channel–opening antibiotic, increases the concentration of cytoplasmic calcium ions (Ca^2+^) in NK cells (42). The activation of PKC forms CARMA/CARD-BCL10-MALT1 (CBM) complexes, and MALT1 controls NF-kB activation (62). In addition, Ca^2+^ influx activates calcineurin and dephosphorylates NFAT1, causing translocation from the cytoplasm into the nucleus (63). Both NF-kB and NFAT1 activate IFN-γ transcription (64). Hence, PMA plus ionomycin can induce IFN-γ production without ligands of NK receptors. Although HLA class 1 molecules and PMA plus ionomycin deploy different mechanisms for IFN-γ production, the amount of IFN-γ was comparable.

Unlike that in NK cells, the production of IFN-γ in KHYG1 cells was dependent on the IL-2 concentration (Figure 5A), despite the surface expression of KLRK1 being higher than that on NK cells (Figure 4B). KHYG1 cells require IL-2 for survival and proliferation (39) and showed lower expression of HCST in KHYG1 cells than in NK cells (Figure 5B), which does not convey HLA-B51 stimulation. Furthermore, the higher expression of IL-2RA in KHYG1 cells than in NK cells (Figure 5D) might strengthen responses to IL-2 because IL-2RA, IL2RB, and IL2RG form receptors with high affinity to IL-2 (40). These findings suggest that HLA-B51 selects higher levels of KLRK1 and promotes IFN-γ production in NK cells (Figure 6A). Pairing of KLRK1 with HCST recruits PI3K and GRB2–VAV1 mediating HLA-B51 stimulation in NK cells (65). In KHYG1 cells, IL-2 signals mediated through IL2Rαβγ override HLA-B51 stimulation and produce IFN-γ (Figure 6B). Furthermore, JAK/STAT could possibly mediate IL-2 signals and induce the production of IFN-γ (66).

**FIGURE 6.**
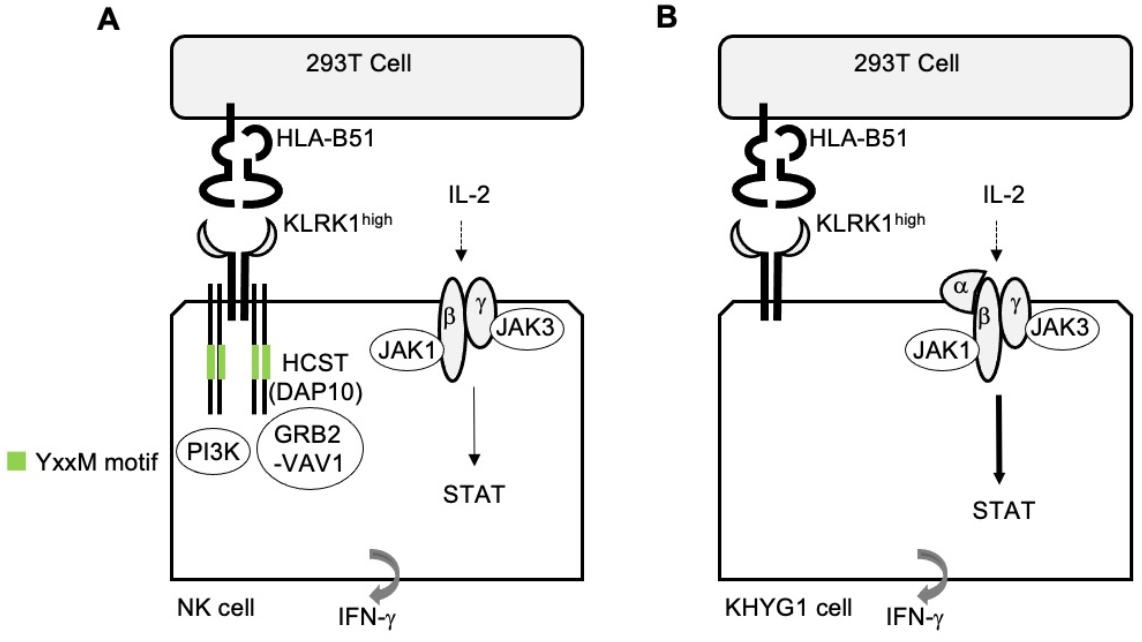
The different mechanisms of IFN-γ production in NK cells and KHYG1 cells (A) HLA-B51 stimulation is mediated through KLRK1 with its adaptor protein HCST and induces IFN-γ secretion from NK cells. (B) In KHYG1 cells, IL2RA binds IL-2 with high affinity and improves IL-2 signaling for IFN-γ production, whereas KLRK1 cannot mediate HLA-B51 stimulation due to low expression of HCST.

## Data availability statement

The original contributions presented in the study are included in the article/Supplementary materials. Further inquiries can be directed to the corresponding author.

## Ethics statement

Peripheral blood was obtained from healthy donors in Jichi Medical University. This study was approved by the Ethics Review Committee for University Clinical Research of Jichi Medical University.

## Author contribution

Y.O. and K.S. designed the study. Y.O. and H. H. contributed to data collection. Y.O. wrote the manuscript. All authors discussed and approved the manuscript.

## Funding

This work was partly supported by the Grants-in-Aid for Scientific Research (C) (No. 23K09156).

## Acknowledgements

We would like to thank for Jun Nakamura for blood collection from healthy donors.

## Conflicts of interest

The authors declare no conflicts of interest.

## Generative AI statement

The authors declare that no Generative AI was used for preparing this manuscript.

## Notes

### Competing Interest Statement

The authors have declared no competing interest.

## References

1. Tugal-Tutkun I, Onal S, Altan-Yaycioglu R, Huseyin Altunbas H, Urgancioglu M. Uveitis in Behcet disease: an analysis of 880 patients. Am J Ophthalmol. (2004) 138(3):373–80. doi: 10.1016/j.ajo.2004.03.022.

2. Tascilar K, Melikoglu M, Ugurlu S, Sut N, Caglar E, Yazici H. Vascular involvement in Behçet’s syndrome: a retrospective analysis of associations and the time course. Rheumatology (Oxford). (2014) 53(11):2018–22. doi: 10.1093/rheumatology/keu233.

3. Emmi G, Bettiol A, Hatemi G, Prisco D. Behcet’s syndrome. Lancet. (2024) 403(10431):1093–108. doi: 10.1016/S0140-6736(23)02629-6.

4. Ohno S, Ohguchi M, Hirose S, Matsuda H, Wakisaka A, Aizawa M. Close association of HLA-Bw51 with Behcet’s disease. Arch Ophthalmol. (1982) 100(9):1455–8. doi: 10.1001/archopht.1982.01030040433013.

5. Mizuki N, Meguro A, Tohnai I, Gül A, Ohno S, Mizuki N. Association of Major Histocompatibility Complex Class I Chain-Related Gene A and HLA-B Alleles with Behçet’s Disease in Turkey. Jpn J Ophthalmol. (2007) 51(6):431–6. doi: 10.1007/s10384-007-0473-y.

6. Remmers EF, Cosan F, Kirino Y, Ombrello MJ, Abaci N, Satorius C, et al. Genome-wide association study identifies variants in the MHC class I, IL10, and IL23R-IL12RB2 regions associated with Behçet’s disease. Nat Genet. (2010) 42(8):698–702. doi: 10.1038/ng.625.

7. Hughes T, Coit P, Adler A, Yilmaz V, Aksu K, Düzgün N, et al. Identification of multiple independent susceptibility loci in the HLA region in Behçet’s disease. Nat Genet. (2013) 45(3):319–24. doi: 10.1038/ng.2551.

8. Ombrello MJ, Kirino Y, de Bakker PI, Gül A, Kastner DL, Remmers EF. Behçet disease-associated MHC class I residues implicate antigen binding and regulation of cell-mediated cytotoxicity. Proc Natl Acad Sci U S A. (2014) 111(24):8867–72. doi: 10.1073/pnas.1406575111.

9. Evans DM, Spencer CC, Pointon JJ, Su Z, Harvey D, Kochan G, et al. Interaction between ERAP1 and HLA-B27 in ankylosing spondylitis implicates peptide handling in the mechanism for HLA-B27 in disease susceptibility. Nat Genet. (2011) 43(8):761–67. doi: 10.1038/ng.873.

10. Robinson PC, Claushuis TA, Cortes A, Martin TM, Evans DM, Leo P, et al. Genetic dissection of acute anterior uveitis reveals similarities and differences in associations observed with ankylosing spondylitis. Arthritis Rheumatol. (2015) 67(1):140–51. doi: 10.1002/art.38873.

11. Kasuya K, Hashimoto Y, Numano F. Left ventricular dysfunction and HLA Bw52 antigen in Takayasu arteritis. Heart Vessels Suppl. (1992) 7:116–9. doi: 10.1007/BF01744556.

12. Futami S, Aoyama N, Honsako Y, Tamura T, Morimoto S, Nakashima T, et al. HLA-DRB1*1502 allele, subtype of DR15, is associated with susceptibility to ulcerative colitis and its progression. Dig Dis Sci. (1995) 40(4):814–18. doi: 10.1007/BF02064985.

13. Allen MH, Ameen H, Veal C, Evans J, Ramrakha-Jones VS, Marsland AM, et al. The major psoriasis susceptibility locus PSORS1 is not a risk factor for late-onset psoriasis. J Invest Dermatol. (2005) 124(1):103–6. doi: 10.1111/j.0022-202X.2004.23511.x.

14. McGonagle D, Aydin SZ, Gül A, Mahr A, Direskeneli H. ‘MHC-I-opathy’-unified concept for spondyloarthritis and Behcet disease. Nat Rev Rheumatol. (2015) 11(12):731–40. doi: 10.1038/nrrheum.2015.147.

15. Ureten K, Ertenli I, Oztürk MA, Kiraz S, Onat AM, Tuncer M, et al. Neutrophil CD64 expression in Behçet’s disease. J Rheumatol. (2005) 32(5):849–52.

16. Becatti M, Emmi G, Silvestri E, Bruschi G, Ciucciarelli L, Squatrito D, et al. Neutrophil Activation Promotes Fibrinogen Oxidation and Thrombus Formation in Behçet Disease. Circulation. (2016) 133(3):302–11. doi: 10.1161/CIRCULATIONAHA.115.017738.

17. Le Joncour A, Martos R, Loyau S, Lelay N, Dossier A, Cazes A, et al. Critical role of neutrophil extracellular traps (NETs) in patients with Behcet’s disease. Ann Rheum Dis. (2019) 78(9):1274–82. doi: 10.1136/annrheumdis-2018-214335.

18. Le Joncour A, Cacoub P, Boulaftali Y, Saadoun D. Neutrophil, NETs and Behcet’s disease: A review. Clin Immunol. (2023) 250:109318. doi: 10.1016/j.clim.2023.109318.

19. van der Houwen TB, van Hagen PM, van Laar JAM. Immunopathogenesis of Behcet’s disease and treatment modalities. Semin Arthritis Rheum. (2022) 52:151956. doi: 10.1016/j.semarthrit.2022.151956.

20. Hasan MS, Bergmeier LA, Petrushkin H, Fortune F. Gamma Delta (γδ) T Cells and Their Involvement in Behçet’s Disease. J Immunol Res. (2015) 2015:705831. doi: 10.1155/2015/705831.

21. Krangel MS, Orr HT, Strominger JL. Assembly and maturation of HLA-A and HLA-B antigens in vivo. Cell. (1979) 18(4):979–91. doi: 10.1016/0092-8674(79)90210-1.

22. Litwin V, Gumperz J, Parham P, Phillips JH, Lanier LL. NKB1: a natural killer cell receptor involved in the recognition of polymorphic HLA-B molecules. J Exp Med. (1994) 180(2):537–43. doi: 10.1084/jem.180.2.537.

23. Pugh J, Nemat-Gorgani N, Djaoud Z, Guethlein LA, Norman PJ, Parham P. In vitro education of human natural killer cells by KIR3DL1. Life Sci Alliance. (2019) 2(6):e201900434.

24. Kim S, Sunwoo JB, Yang L, Choi T, Song YJ, French AR, et al. HLA alleles determine differences in human natural killer cell responsiveness and potency. Proc Natl Acad Sci U S A. (2008) 105(8):3053–8. doi: 10.1073/pnas.0712229105.

25. Hasan MS, Ryan PL, Bergmeier LA, Fortune F. Circulating NK cells and their subsets in Behçet’s disease. Clin Exp Immunol. (2017) 188(2):311–22. doi: 10.1111/cei.12939.

26. Deniz G, Akdis M, Aktas E, Blaser K, Akdis CA. Human NK1 and NK2 subsets determined by purification of IFN-gamma-secreting and IFN-gamma-nonsecreting NK cells. Eur J Immunol. (2002) 32(3):879–84. doi: 10.1002/1521-4141(200203)32:3<879::AID-IMMU879>3.0.CO;2-2.

27. Kucuksezer UC, Aktas-Cetin E, Bilgic-Gazioglu S, Tugal-Tutkun I, Gül A, Deniz G. Natural killer cells dominate a Th-1 polarized response in Behcet’s disease patients with uveitis. Clin Exp Rheumatol. (2015) 33(6 Suppl 94):S24–9.

28. Cosan F, Aktas Cetin E, Akdeniz N, Emrence Z, Cefle A, Deniz G. Natural Killer Cell Subsets and Their Functional Activity in Behçet’s Disease. Immunol Invest. (2017) 46(4):419–32. doi: 10.1080/08820139.2017.1288240.

29. Zheng W, Wang X, Liu J, Yu X, Li L, Wang H, et al. Single-cell analyses highlight the proinflammatory contribution of C1q-high monocytes to Behçet’s disease. Proc Natl Acad Sci U S A. (2022) 119(26):e2204289119. doi: 10.1073/pnas.2204289119.

30. Wu J, Song Y, Bakker AB, Bauer S, Spies T, Lanier LL, et al. An activating immunoreceptor complex formed by NKG2D and DAP10. Science. (1999) 285(5428):730–32. doi: 10.1126/science.285.5428.730.

31. Billadeau DD, Upshaw JL, Schoon RA, Dick CJ, Leibson PJ. NKG2D-DAP10 triggers human NK cell-mediated killing via a Syk-independent regulatory pathway. Nat Immunol. (2003) 4(6):557–64. doi: 10.1038/ni929.

32. Bauer S, Groh V, Wu J, Steinle A, Phillips JH, Lanier LL, et al. Activation of NK cells and T cells by NKG2D, a receptor for stress-inducible MICA. Science. (1999) 285(5428):727–29. doi: 10.1126/science.285.5428.727.

33. Cosman D, Müllberg J, Sutherland CL, Chin W, Armitage R, Fanslow W, et al. ULBPs, novel MHC class I-related molecules, bind to CMV glycoprotein UL16 and stimulate NK cytotoxicity through the NKG2D receptor. Immunity. (2001) 14(2):123–33. doi: 10.1016/s1074-7613(01)00095-4.

34. Sutherland CL, Chalupny NJ, Schooley K, VandenBos T, Kubin M, Cosman D. UL16-binding proteins, novel MHC class I-related proteins, bind to NKG2D and activate multiple signaling pathways in primary NK cells. J Immunol. (2002) 168(2):671–9. doi: 10.4049/jimmunol.168.2.671.

35. Sivori S, Pende D, Bottino C, Marcenaro E, Pessino A, Biassoni R, et al. NKp46 is the major triggering receptor involved in the natural cytotoxicity of fresh or cultured human NK cells. Correlation between surface density of NKp46 and natural cytotoxicity against autologous, allogeneic or xenogeneic target cells. Eur J Immunol. (1999) 29(5):1656–66. doi: 10.1002/(SICI)1521-4141(199905)29:05<1656::AID-IMMU1656>3.0.CO;2-1.

36. Omata Y, Tachibana H, Aizaki Y, Mimura T, Sato K. Essentiality of Nfatc1 short isoform in osteoclast differentiation and its self-regulation. Sci Rep. (2023) 13(1):18797. doi: 10.1038/s41598-023-45909-3.

37. Jones DC, Edgar RS, Ahmad T, Cummings JR, Jewell DP, Trowsdale J, et al. Killer Ig-like receptor (KIR) genotype and HLA ligand combinations in ulcerative colitis susceptibility. Genes Immun. (2006) 7(7):576–82. doi: 10.1038/sj.gene.6364333.

38. Dellgren C, Nehlin JO, Barington T. Cell surface expression level variation between two common Human Leukocyte Antigen alleles, HLA-A2 and HLA-B8, is dependent on the structure of the C terminal part of the alpha 2 and the alpha 3 domains. PLoS One. (2015) 10(8):e0135385. doi: 10.1371/journal.pone.0135385.

39. Yagita M, Huang CL, Umehara H, Matsuo Y, Tabata R, Miyake M, et al. A novel natural killer cell line (KHYG-1) from a patient with aggressive natural killer cell leukemia carrying a p53 point mutation. Leukemia. (2000) 14(5):922–30. doi: 10.1038/sj.leu.2401769.

40. Kim HP, Imbert J, Leonard WJ. Both integrated and differential regulation of components of the IL-2/IL-2 receptor system. Cytokine Growth Factor Rev. (2006) 17(5):349–66. doi: 10.1016/j.cytogfr.2006.07.003.

41. Kaszubowska L, Foerster J, Schetz D, Kmieć Z. CD56bright cells respond to stimulation until very advanced age revealing increased expression of cellular protective proteins SIRT1, HSP70 and SOD2. Immun Ageing. (2018) 15:31. doi: 10.1186/s12979-018-0136-5.

42. Adib Y, Boy M, Serror K, Dulphy N, des Courtils C, Duciel L, et al. Modulation of NK cell activation by exogenous calcium from alginate dressings in vitro. Front Immunol. (2023) 14:1141047. doi: 10.3389/fimmu.2023.1141047.

43. Wulff S, Pries R, Wollenberg B. Cytokine release of human NK cells solely triggered with Poly I:C. Cell Immunol. (2010) 263(2):135–37. doi: 10.1016/j.cellimm.2010.03.020.

44. Rajagopalan S, Fu J, Long EO. Cutting edge: induction of IFN-gamma production but not cytotoxicity by the killer cell Ig-like receptor KIR2DL4 (CD158d) in resting NK cells. J Immunol. (2001) 167(4):1877–81. doi: 10.4049/jimmunol.167.4.1877.

45. Kikuchi-Maki A, Yusa S, Catina TL, Campbell KS. KIR2DL4 is an IL-2-regulated NK cell receptor that exhibits limited expression in humans but triggers strong IFN-gamma production. J Immunol. (2003) 171(7):3415–25. doi: 10.4049/jimmunol.171.7.3415.

46. Fauriat C, Long EO, Ljunggren HG, Bryceson YT. Regulation of human NK-cell cytokine and chemokine production by target cell recognition. Blood. (2010) 115(11):2167–76. doi: 10.1182/blood-2009-08-238469.

47. Hammond ME, Lapointe GR, Feucht PH, Hilt S, Gallegos CA, Gordon CA, et al. IL-8 induces neutrophil chemotaxis predominantly via type I IL-8 receptors. J Immunol. (1995) 155(3):1428–33.

48. Shu Q, Zhang N, Liu Y, Wang X, Chen J, Xie H, et al. IL-8 Triggers Neutrophil Extracellular Trap Formation Through an Nicotinamide Adenine Dinucleotide Phosphate Oxidase- and Mitogen-Activated Protein Kinase Pathway-Dependent Mechanism in Uveitis. Invest Ophthalmol Vis Sci. (2023) 64(13):19. doi: 10.1167/iovs.64.13.19.

49. Durmazlar SP, Ulkar GB, Eskioglu F, Tatlican S, Mert A, Akgul A. Significance of serum interleukin-8 levels in patients with Behcet’s disease: high levels may indicate vascular involvement. Int J Dermatol. (2009) 48(3):259–64. doi: 10.1111/j.1365-4632.2009.03905.x.

50. Young JD, Hengartner H, Podack ER, Cohn ZA. Purification and characterization of a cytolytic pore-forming protein from granules of cloned lymphocytes with natural killer activity. Cell. (1986) 44(6):849–59. doi: 10.1016/0092-8674(86)90007-3.

51. Su B, Bochan MR, Hanna WL, Froelich CJ, Brahmi Z. Human granzyme B is essential for DNA fragmentation of susceptible target cells. Eur J Immunol. (1994) 24(9):2073–80. doi: 10.1002/eji.1830240921.

52. Reefman E, Kay JG, Wood SM, Offenhäuser C, Brown DL, Roy S, et al. Cytokine secretion is distinct from secretion of cytotoxic granules in NK cells. J Immunol. (2010) 184(9):4852–62. doi: 10.4049/jimmunol.0803954.

53. Tóth J and Kubes M. Masking of HLA class I molecules expressed on K-562 target cells can restore their susceptibility to NK cell cytolysis. Immunobiology. (1993) 188(1–2):134–44. doi: 10.1016/S0171-2985(11)80493-6.

54. Fujisaki H, Kakuda H, Shimasaki N, Imai C, Ma J, Lockey T, et al. Expansion of highly cytotoxic human natural killer cells for cancer cell therapy. Cancer Res. (2009) 69(9):4010–17. doi: 10.1158/0008-5472.CAN-08-3712.

55. Lanier LL, Corliss B, Wu J, Phillips JH. Association of DAP12 with activating CD94/NKG2C NK cell receptors. Immunity. (1998) 8(6):693–701. doi: 10.1016/s1074-7613(00)80574-9.

56. Moretta A, Vitale M, Sivori S, Bottino C, Morelli L, Augugliaro R, et al. Human natural killer cell receptors for HLA-class I molecules. Evidence that the Kp43 (CD94) molecule functions as receptor for HLA-B alleles. J Exp Med. (1994) 180(2):545–55. doi: 10.1084/jem.180.2.545.

57. Karimi MA, Aguilar O, Zou B, Bachmann MH, Carlyle JR, Baldwin CL, et al. A truncated human NKG2D splice isoform negatively regulates NKG2D-mediated function. J Immunol. (2014) 193(6):2764–71. doi: 10.4049/jimmunol.1400920.

58. Plougastel B and Trowsdale J. Cloning of NKG2-F, a new member of the NKG2 family of human natural killer cell receptor genes. Eur J Immunol. (1997) 27(11):2835–39. doi: 10.1002/eji.1830271114.

59. Kim DK, Kabat J, Borrego F, Sanni TB, You CH, Coligan JE. Human NKG2F is expressed and can associate with DAP12. Mol Immunol. (2004) 41(1):53–62. doi: 10.4049/jimmunol.174.5.2878.

60. Caldas P, Luz M, Baseggio S, Andrade R, Sobral D, Grosso AR. Transcription readthrough is prevalent in healthy human tissues and associated with inherent genomic features. Commun Biol. (2024) 7(1):100. doi: 10.1038/s42003-024-05779-5.

61. Cai G, Kastelein RA, Hunter CA. IL-10 enhances NK cell proliferation, cytotoxicity and production of IFN-gamma when combined with IL-18. Eur J Immunol. (1999) 29(9):2658–65. doi: 10.1002/(SICI)1521-4141(199909)29:09<2658::AID-IMMU2658>3.0.CO;2-G.

62. Juilland M and Thome M. Holding All the CARDs: How MALT1 Controls CARMA/CARD-Dependent Signaling. Front Immunol. (2018) 9:1927. doi: 10.3389/fimmu.2018.01927. eCollection 2018.

63. Hogan PG, Chen L, Nardone J, Rao A. Transcriptional regulation by calcium, calcineurin, and NFAT. Genes Dev. (2003) 17(18):2205–32. doi: 10.1101/gad.1102703.

64. Sica A, Dorman L, Viggiano V, Cippitelli M, Ghosh P, Rice N, et al, Interaction of NF-kappaB and NFAT with the interferon-gamma promoter. J Biol Chem. (1997) 272(48):30412–20. doi: 10.1074/jbc.272.48.30412.

65. Wilton KM, Overlee BL, Billadeau DD. NKG2D-DAP10 signaling recruits EVL to the cytotoxic synapse to generate F-actin and promote NK cell cytotoxicity. J Cell Sci. (2019) 133(5):jcs230508.

66. Gotthardt D, Trifinopoulos J, Sexl V, Putz EM. JAK/STAT Cytokine Signaling at the Crossroad of NK Cell Development and Maturation. Front Immunol. (2019) 10:2590. doi: 10.3389/fimmu.2019.02590.

