## Supplementary Figures for "HLA-B51 induces IFN-*γ* production in human natural killer cells": 260502bn_2 omata y et al supplementary material.docx

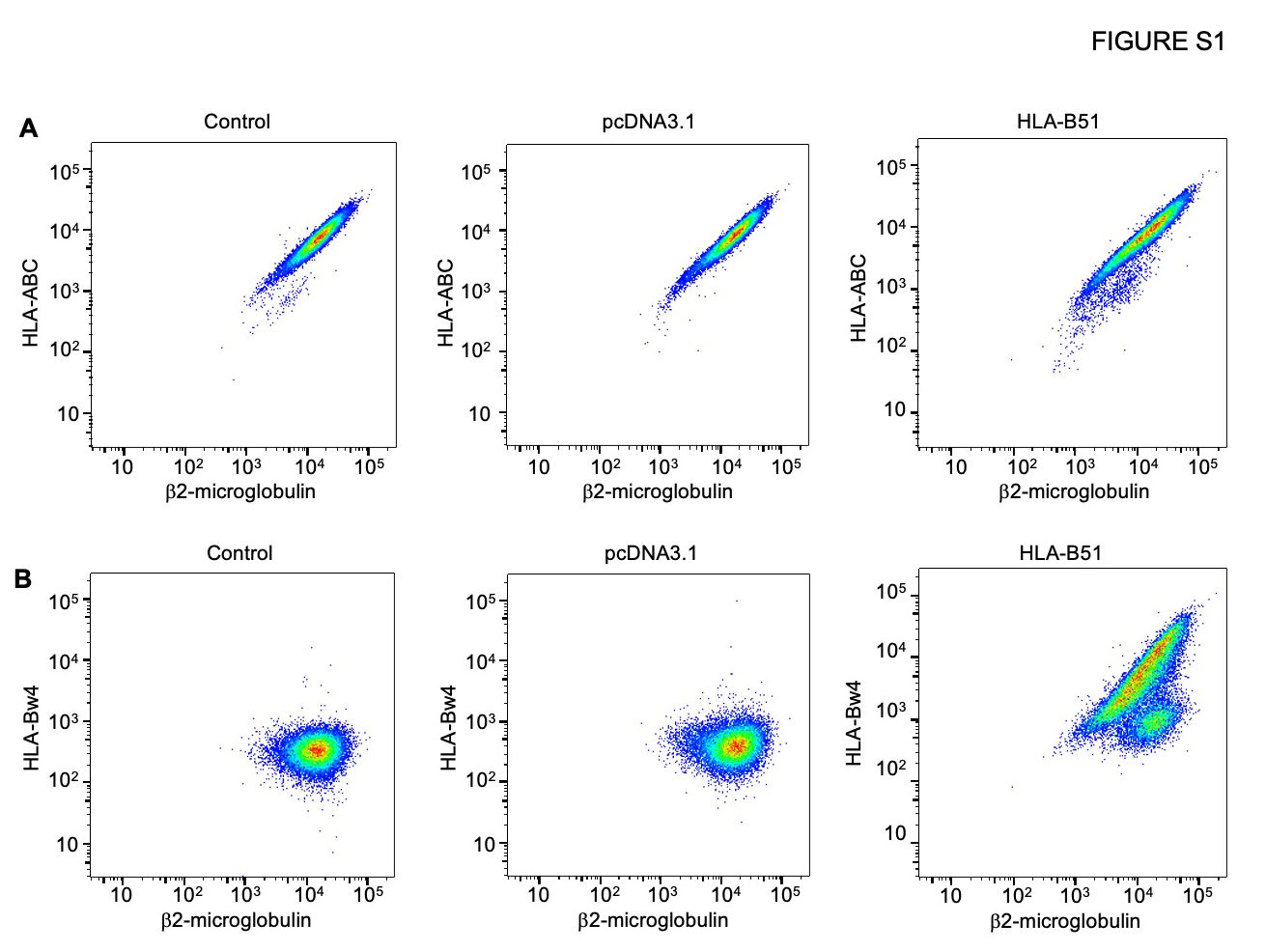

**FIGURE S1 HLA class 1 complex on 293T cells.**

(A, B) Plasmid DNA was transfected into 293T cells. At 24 h, the cell surface expression in 293T cells was analyzed by flow cytometry using anti-HLA-ABC, anti-HLA-Bw4, and anti-β2-microglobulin antibodies. Representative data of the surface expression of β2-microglobulin with HLA-ABC (A) and HLA-Bw4 (B) are shown.

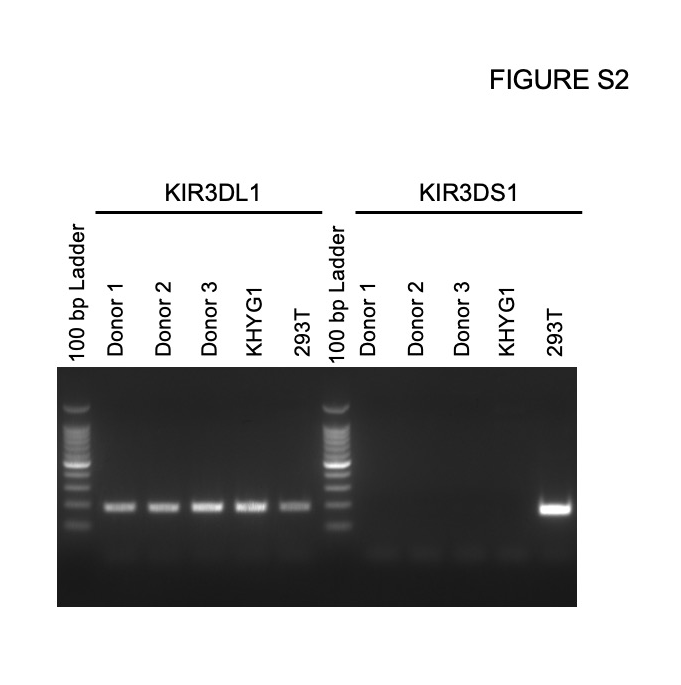

**FIGURE S2 KIR typing by PCR.**

Genomic DNA was extracted from NK cells, KHYG1 cells, and 293T cells. KIR3DL1 and KLR3DS1 types were tested by PCR.
